# L-DOPA treatment promotes sustained neurovascular and synaptic homeostasis in the diabetic retina

**DOI:** 10.64898/2026.02.17.706466

**Authors:** Eli Chlan, Chenxing Li, Katie L. Bales, Levi B. Wood, Machelle T. Pardue

## Abstract

While previous work has shown a sustained protective effect of levodopa (L-DOPA) on retinal function in early-stage diabetic retinopathy (DR) in humans, its underlying biology is unknown. Using noninvasive measures in diabetic mice, we found L-DOPA protects retinal neurovascular function as measured by oscillatory potential timing and flicker-evoked retinal vasodilation, as well as visual behavior, for at least two weeks past treatment end. Assessing changes in retinal gene expression, differentially expressed genes were broadly comparable between diabetic mice experiencing washout of L-DOPA versus continued L-DOPA treatment, with gene co-expression network analysis identifying distinct modules across L-DOPA–treated diabetic mice associated with synaptic function and cytoskeletal organization that correlated with functional protection. Together, these findings demonstrate that L-DOPA restores and sustains retinal neurovascular function in early DR and links this protection to transcriptional programs supporting synapse activity and structural integrity.

**Teaser:** L-DOPA restores neuronal and vascular performance in the diabetic retina, with benefits that lasted after treatment stopped.

## Introduction

Diabetic retinopathy (DR) has historically been defined as a retinal microvascular complication resulting from diabetes mellitus (DM) and has more recently been identified as a neurovascular disease (*1–3*). One-third of diabetic patients develop DR, with a further third of DR patients experiencing vision-threatening DR (*4*). Clinical detection and classification of DR relies on visualizing the development of vascular manifestations, such as microaneurysms, venous beading, retinal and vitreous hemorrhages, gliosis and/or retinal neovascularization (*2, 5, 6*). While convenient for diagnosis, ongoing research has revealed neuronal and vascular dysfunction that precede these large-scale, vision-threatening vascular changes (*7–12*).

These early deficits in the diabetic retina span the neurovascular unit (NVU) - an intricate coupling of neurons, glia, immune cells, and vasculature within the central nervous system. Unlike the rest of the eye, the retina lacks autonomic innervation and relies on the NVU for autoregulation of retinal circulation (*13*). In preclinical DR, changes across cell types of the NVU include delay in rod-specific neuronal signaling (*7–11*), reduction of neuronal synaptic connectivity (*12, 14*), gliosis by Müller cells (*12*), reduction of blood flow and basal vessel diameter (*15–17*), and increasing interactions of glia (e.g. microglia) with retinal vasculature (*18*). Several studies show that individual deficits within the NVU of the diabetic retina disrupt the function of its associated components (*19–21*). Given this interdependence, identifying treatments for DR that target these early functional deficits in the retina may be critical to deterring progression towards vision-threatening vascular complications.

In the retina, dopamine is a critical neuromodulator that shapes gap junction coupling and intrinsic ionic conductance, facilitating dynamic modulation of retinal circuitry to optimize performance under different lighting conditions (*22–24*). Retinal dopamine levels are decreased in diabetic rodent models, with reduced dopamine concomitant with early functional changes, such as amacrine cell-derived oscillatory potential implicit timing (OP IT) delays (*8, 25*). Importantly, restoring dopamine in diabetic rodent models by administration of dopamine’s precursor, levodopa (L-DOPA), was protective against OP IT delays and visual function deficit when given at hyperglycemic onset (*8, 11*), or administered after OP IT delays were detected (*9*). Strikingly, translation to diabetic patients showed that L-DOPA was restorative for OP IT delays and that this neuroprotective effect was sustained for at least two weeks beyond the treatment window (*10*).

However, biological underpinnings relevant to L-DOPA’s sustained functional neuroprotection in the diabetic retina remain unexplored. Here, we show L-DOPA benefits retinal neurovascular coupling and replicate L-DOPA’s lasting protective effects on retinal function in diabetic mice. Further, we establish L-DOPA-induced transcriptional changes in the diabetic retina, with gene networks spanning synapse signaling and cytoskeletal components correlative with neurovascular functional protection sustained after L-DOPA washout.

## Results

### Sustained benefit of L-DOPA treatment for rod-driven inner retinal function in diabetic mice

Prioritizing clinical relevance, L-DOPA or vehicle treatments were administered to all groups only after retinal and visual functional deficits were detected in hyperglycemic mice (**Fig. 1**). Hyperglycemia (>250 mg/dL) developed in STZ-administered mice after one week and was confirmed across longitudinal measures (**fig. S1, A and B**). After, on average, 7.33±0.67 (mean±SEM) weeks of hyperglycemia, diabetic mice developed significantly delayed oscillatory potential (OP) implicit timing (IT; **Fig. 2A;** Ctrl: 61.5±0.98 ms vs. DM: 68.8±1.03 ms, p<0.0001).

**Fig. 1.**
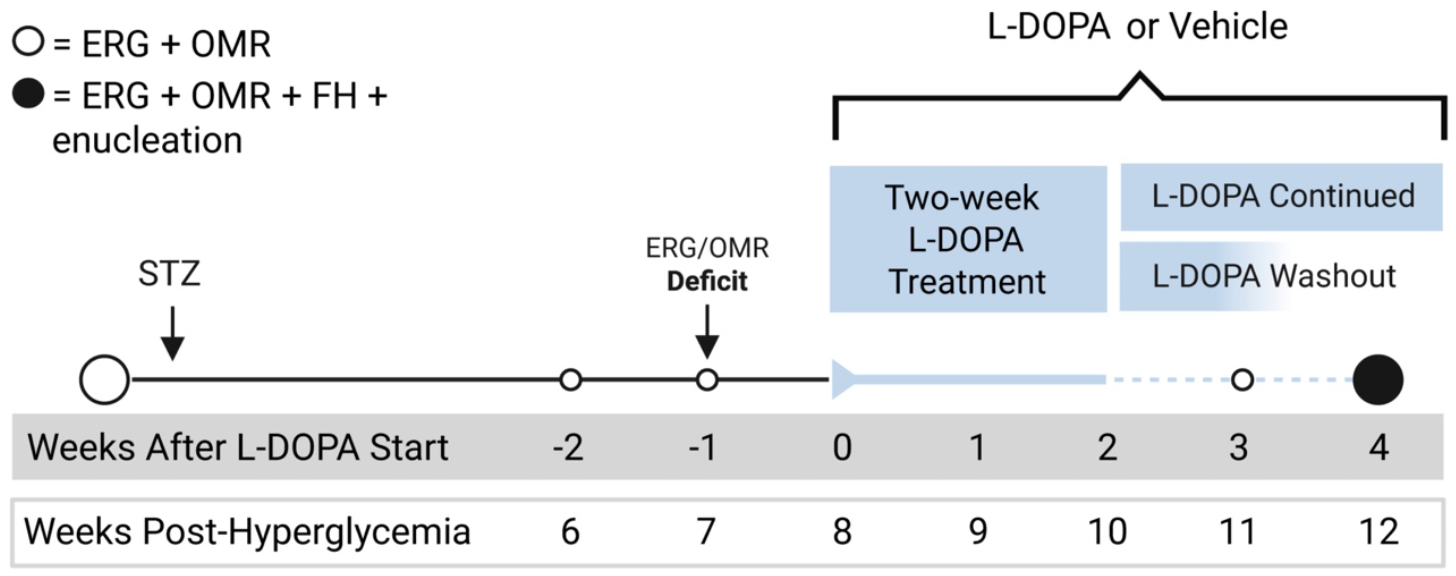
Timeline of experimental design for L-DOPA treatment in the diabetic mouse. Baseline measurements [electroretinograms (ERGs), optomotor response (OMRs)] were taken before mice were made diabetic via STZ or kept as controls. After ERG/OMR deficits were detected, mice received two-week L-DOPA or vehicle treatment, followed by two weeks of either continued L-DOPA/vehicle treatment or no treatment (“washout”). At the final timepoint, ERGs, OMRs and functional hyperemia (FH) were measured and followed by enucleation.

**Fig. 2.**
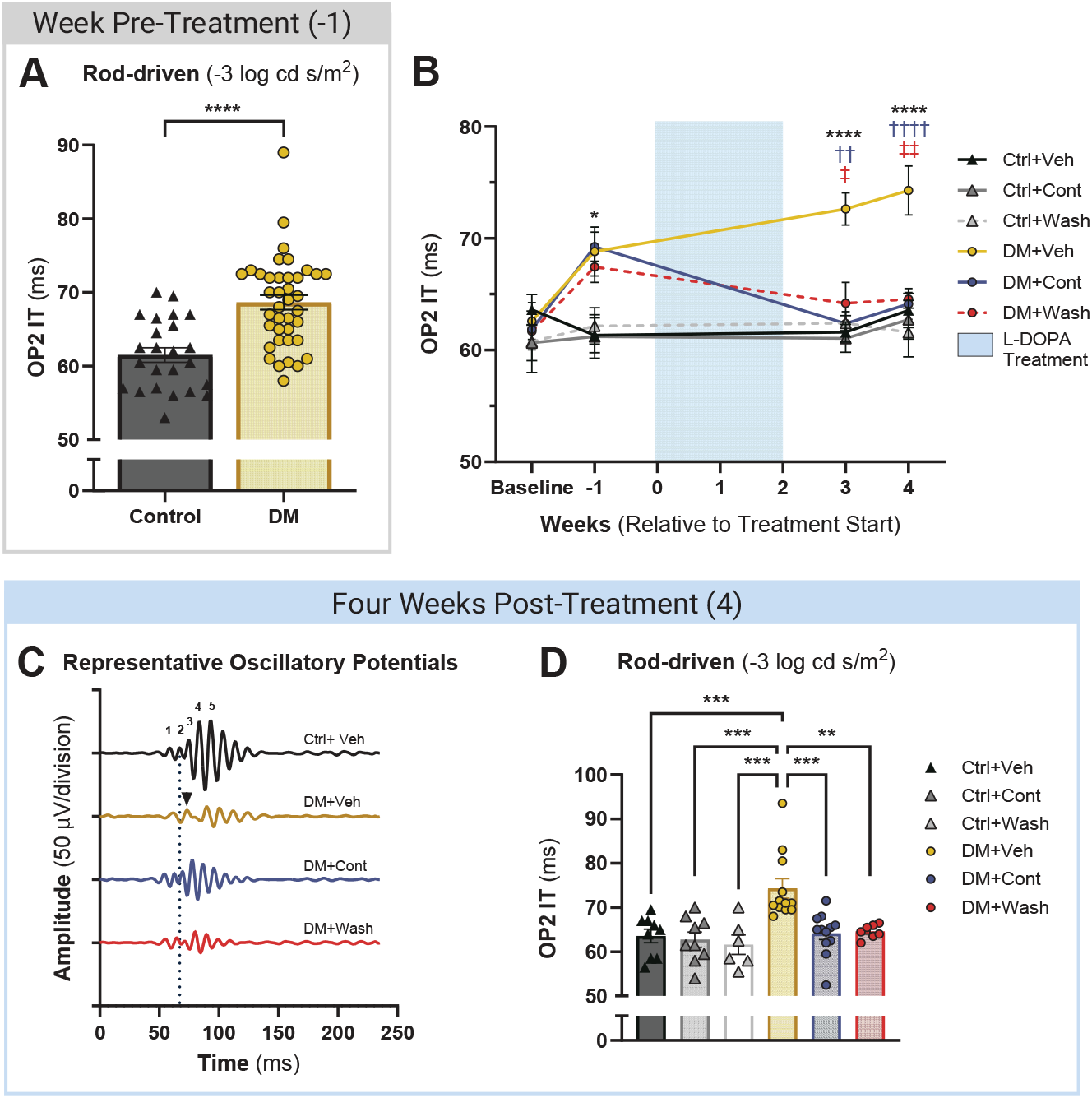
L-DOPA sustained improvement of rod-driven oscillatory potential (OP) timing. (**A**) Rod-driven OP timing (OP2 IT) was significantly delayed in diabetic mice compared to control mice prior to L-DOPA treatment. (**B**) After two weeks of L-DOPA treatment, DM+Cont and DM+Wash mice consistently showed improved OP2 IT over DM+Veh mice to the final timepoint, while DM+Veh OP2 IT grew significantly more delayed. Ctrl+Cont/Wash comparison statistics are not shown here. (**C**) Representative OP waveforms from individual mice at the final experimental timepoint show DM+Veh OP2 IT delay (black arrowhead) versus the timing of Ctrl+Veh OP2 IT (vertical dotted line). Note OP2 is similar or faster in the Cont and Wash L-DOPA groups. (**D**) Prior to enucleation, OP2 IT was similar between DM+Cont, DM+Wash and control mice, while significantly delayed in DM+Veh mice compared to all other groups. Data shown as mean ± SEM; (*) DM+Veh vs. Ctrl+Veh, (†) DM+Veh vs. DM+Cont, (‡) DM+Veh vs. DM+Wash; \*\**p* < 0.01,\*\*\**p* < 0.001,\*\*\*\**p* < 0.0001.

After four weeks of continuous L-DOPA treatment following delay detection, DM+Cont mice had significantly improved OP IT compared to DM+Veh (p<0.0001; **Fig. 2B-D, Table 1**) and were statistically indistinguishable from Ctrl+Veh OP2 IT (p=0.999; **Fig. 2B-D**). DM mice treated with L-DOPA for two weeks followed by two weeks of washout (DM+Wash) also reflected control-like timing and maintained improved OP2 IT over DM+Veh (p=0.004). Similar to our previous findings (*8, 11*), improvement in OP timing in DM+Cont and DM+Wash mice was exclusively rod-driven, with no significant differences in OP IT elicited by brighter flash stimuli (**fig. S2, A**). Furthermore, non-DM Ctrl mice receiving L-DOPA under continued (Ctrl+Cont) or washout (Ctrl+Wash) conditions showed no change to amplitude or timing across OPs, B-wave, and A-wave (**Fig. 2B-D**; **fig. S2**).

**Table 1.**
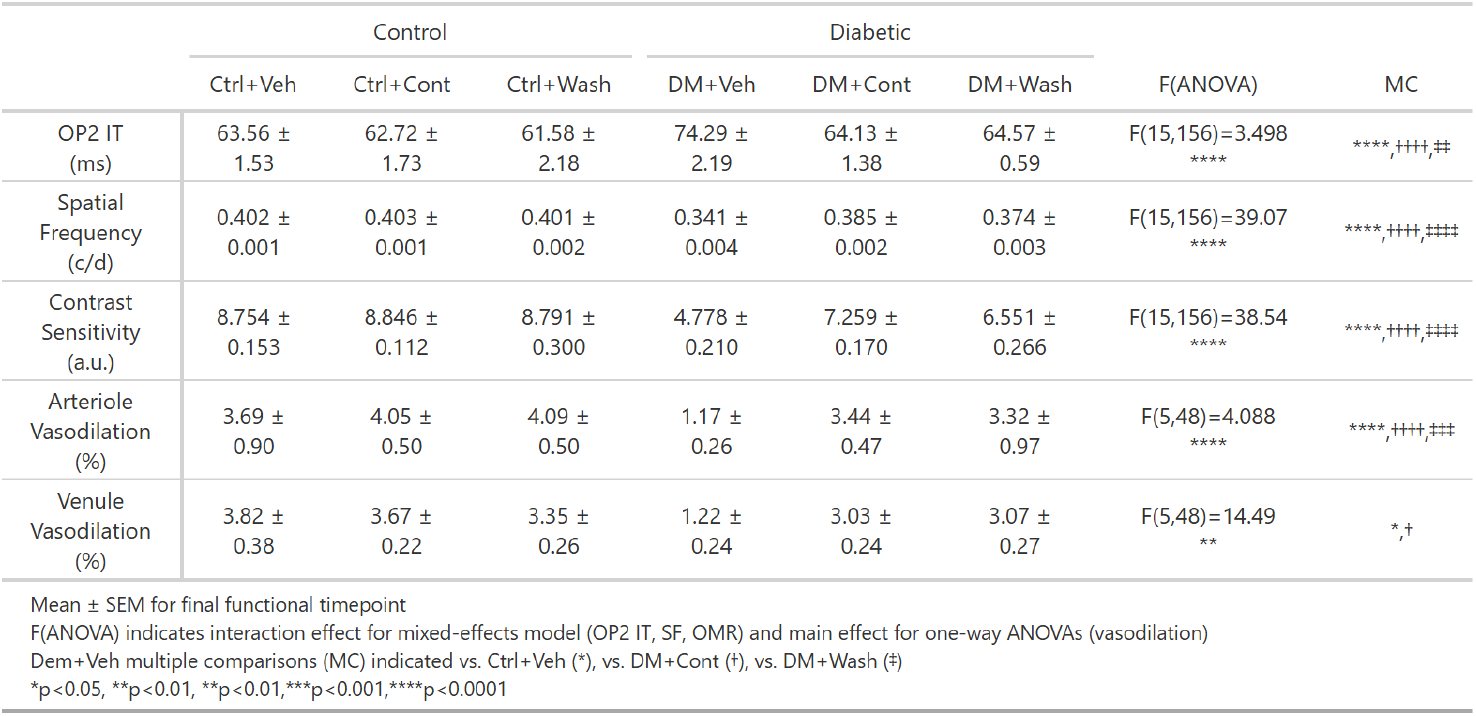
*In vivo* functional assessment from the final timepoint (4 weeks post-treatment) assessed across DM condition and L-DOPA treatment groups.

### L-DOPA induced lasting visual function protection in the diabetic retina

Similar to retinal function deficit detected by ERGs, DM mice had reduced spatial frequency (SF; Ctrl: 0.402±0.0006 c/d vs. DM: 0.367±0.002 c/d, p<0.0001) and contrast sensitivity (CS; Ctrl: 8.66±0.075 a.u. vs. DM: 5.84±0.068 a.u., p<0.0001) thresholds by seven weeks of hyperglycemia (**Fig. 3A**). In DM+Cont and DM+Wash mice, the decline in SF thresholds was halted at three weeks (p<0.0001) and four weeks of treatment (p<0.0001) while DM+Veh SF steadily worsened (**Fig. 3A**; **Table 1**). A similarly robust effect was seen in CS thresholds, with DM+Cont and DM+Wash mice having higher CS thresholds than DM+Veh at three weeks (p<0.0001) and four weeks (p<0.0001) of L-DOPA treatment (**Fig. 3B, Table 1**). While SF and CS were significantly protected from decline, DM+Cont mice had higher thresholds than DM+Wash for SF (p=0.0248) and trending for CS (p=0.0551) at the final timepoint, possibly indicating the start of diverging visual function protection from L-DOPA treatment.

**Fig. 3.**
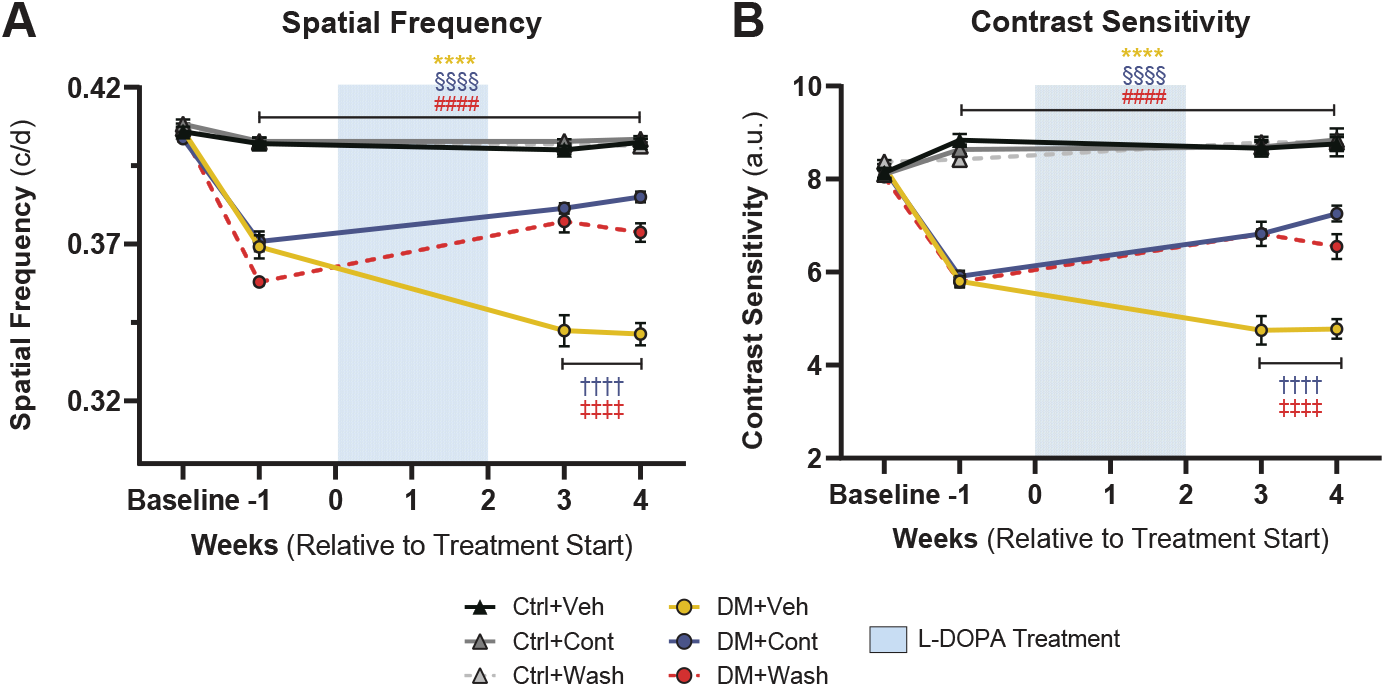
Visual function improvement sustained with L-DOPA treatment in diabetic mice. (**A**) Diabetic mice showed significant reductions in spatial frequency (SF) thresholds prior to L-DOPA treatment (-1 wk treatment). Following two weeks of L-DOPA treatment, DM+Cont and DM+Wash mice demonstrated significantly improved SF thresholds compared to DM+Veh, with dysfunction plateauing compared to continued SF decline in DM+Veh. (**B**) As seen in SF, contrast sensitivity (CS) thresholds significantly declined in diabetic mice before treatment started. L-DOPA treatment induced significantly improved CS thresholds and a similar plateauing effect in DM+Cont and DM+Wash mice compared to DM+Veh mice. Data shown as mean ± SEM; (*) DM+Veh vs. Ctrl+Veh, (†) DM+Veh vs. DM+Cont, (‡) DM+Veh vs. DM+Wash, (§) Ctrl+Veh vs. DM+Cont, (#) Ctrl+Veh vs. DM+Wash;\*\*\*\**p* < 0.0001.

### L-DOPA treatment maintained retinal vascular function in the diabetic retina

To assess the integrity of neurovascular coupling in the diabetic retina, the vasodilative response to increased neuronal demand elicited by 12 Hz photic stimulus was measured across retinal arterioles and venules using confocal scanning laser ophthalmoscopy (**Fig. 4A, B**). In response to light stimulus, Ctrl+Veh arterioles vasodilated 3.68±0.89% while venules vasodilated 3.82± 0.38% from baseline vessel caliber (**Fig. 4C, D**). Arteriole vasodilation values matched flicker-induced observations in clinical (*15, 26*) and murine experiments (*27, 28*). In addition, venule vasodilation mirrored recent reports of measurable venule dilative response by rodents downstream of functional hyperemia activity (*28–30*). As established in previous literature (*27*), DM+Veh light-evoked vasodilation decreased compared to Ctrl+Veh, by 68% in venules (p<0.0001; **Fig. 4C**) and by 68.2% in arterioles (p=0.0323; **Fig. 4D**) after 12 weeks of hyperglycemia.

**Fig. 4.**
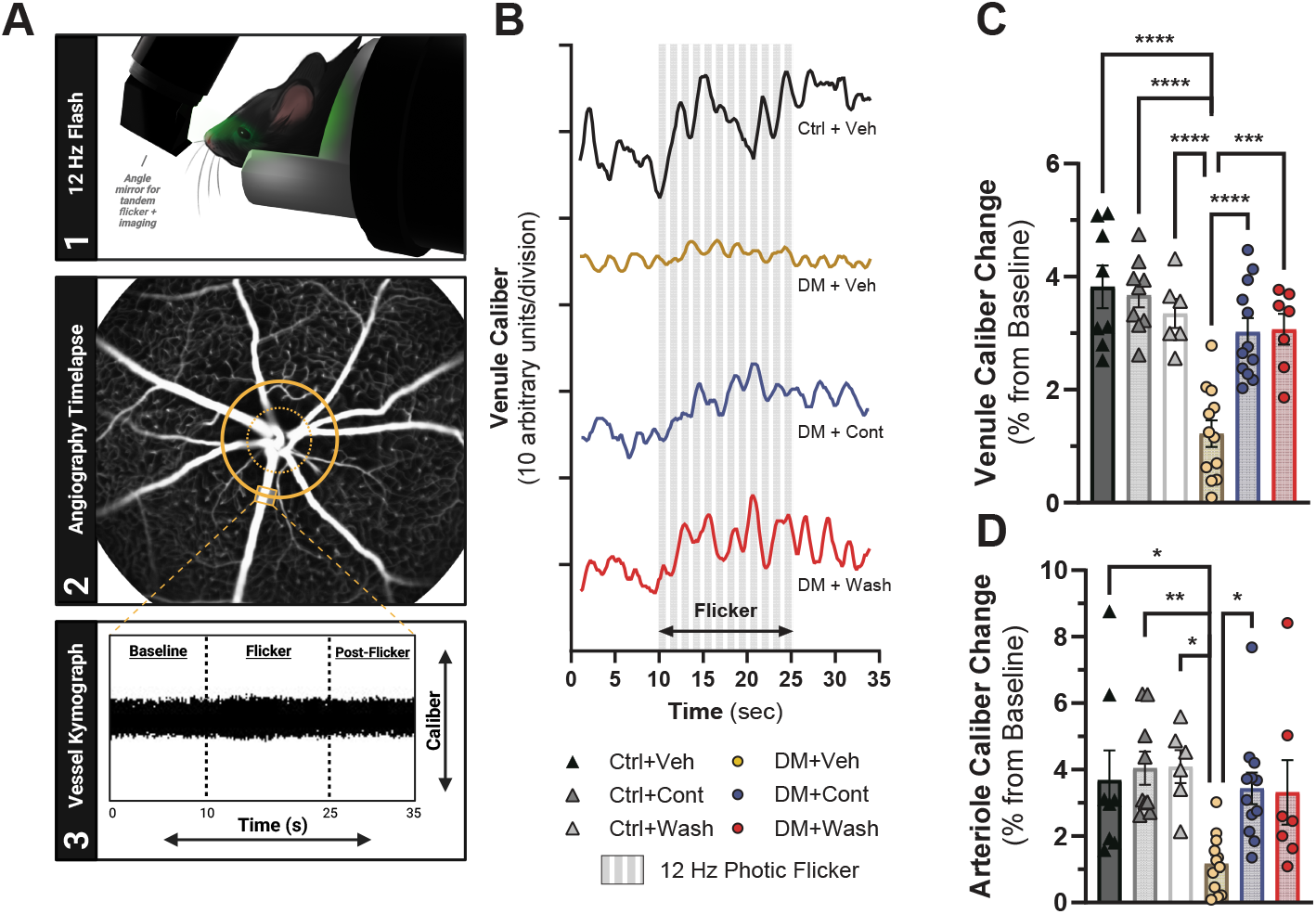
Flicker-induced retinal vasodilation improved with L-DOPA treatment in diabetic mice. (**A**) Graphical representation of *in vivo* neurovascular coupling imaging and analysis pipeline shows (A1) the set-up for stimulating the eye, (A2) the ICG-filled retinal blood vessels visualized with confocal scanning laser ophthalmoscopy, and (A3) a kymograph illustrating vessel dilation across time. (**B**) Representative individual venule caliber traces across pre-flicker, 15-second flicker (gray bars), and post-flicker period. (**C**) Averaged venule caliber percent change from baseline was significantly reduced in DM+Veh compared to control groups. Meanwhile, DM+Cont and DM+Wash mice showed significantly elevated venule response over DM+Veh. (**D**) Similarly, DM+Veh mice had significantly reduced arteriole caliber change compared to control groups. While both DM+Cont and DM+Wash showed elevated arteriole caliber change over DM+Veh, only DM+Cont reached statistical significance. Data shown as mean ± SEM; \**p* < 0.05, \*\**p* < 0.01, \*\**p* < 0.01,\*\*\**p* < 0.001,\*\*\*\**p* < 0.0001.

After four weeks of continuous L-DOPA treatment, DM+Cont mice showed improvement over DM+Veh mice in both arteriole (p=0.0307) and venule (p<0.001) vasodilation (**Fig. 4B-D, Table 1**). Strikingly, following two weeks of L-DOPA treatment washout, DM+Wash mice also had sustained vascular protection as shown by greater venule vasodilation post-washout compared to DM+Veh (p=0.0002; **Fig. 4B, C**; **Table 1**).

Arteriole vasodilation showed similar trends for DM+Wash, but did not reach statistical significance compared to DM+Veh (p=0.124; **Fig. 4D, Table 1**).

### L-DOPA induces lasting synapse-related transcriptional change in diabetic mice

To identify molecular changes driven by L-DOPA’s sustained protective effects, transcriptional profiles across treatment groups were assessed in whole retinal tissue by bulk RNASeq and analyzed for correlation with functional measures. Among 13,665 genes retained after filtering, 489 differentially expressed genes (DEGs) (229 downregulated, 260 upregulated: **Fig. 5A**) were identified in DM-Veh whole retina compared to Ctrl-Veh, outlining transcriptional changes due to the DM model. With L-DOPA treatment, 118 DEGs (10 downregulated, 108 upregulated) were identified in DM-Cont retina compared to DM-Veh, while only 57 DEGs (38 upregulated, 19 downregulated) were identified in DM-Wash compared to DM-Veh (**Fig. 5A**). Over 50% of DM+Wash DEGs identified were shared with DM-Cont DEGs and maintained the same direction of regulation (**Fig. 5B**).

**Fig. 5.**
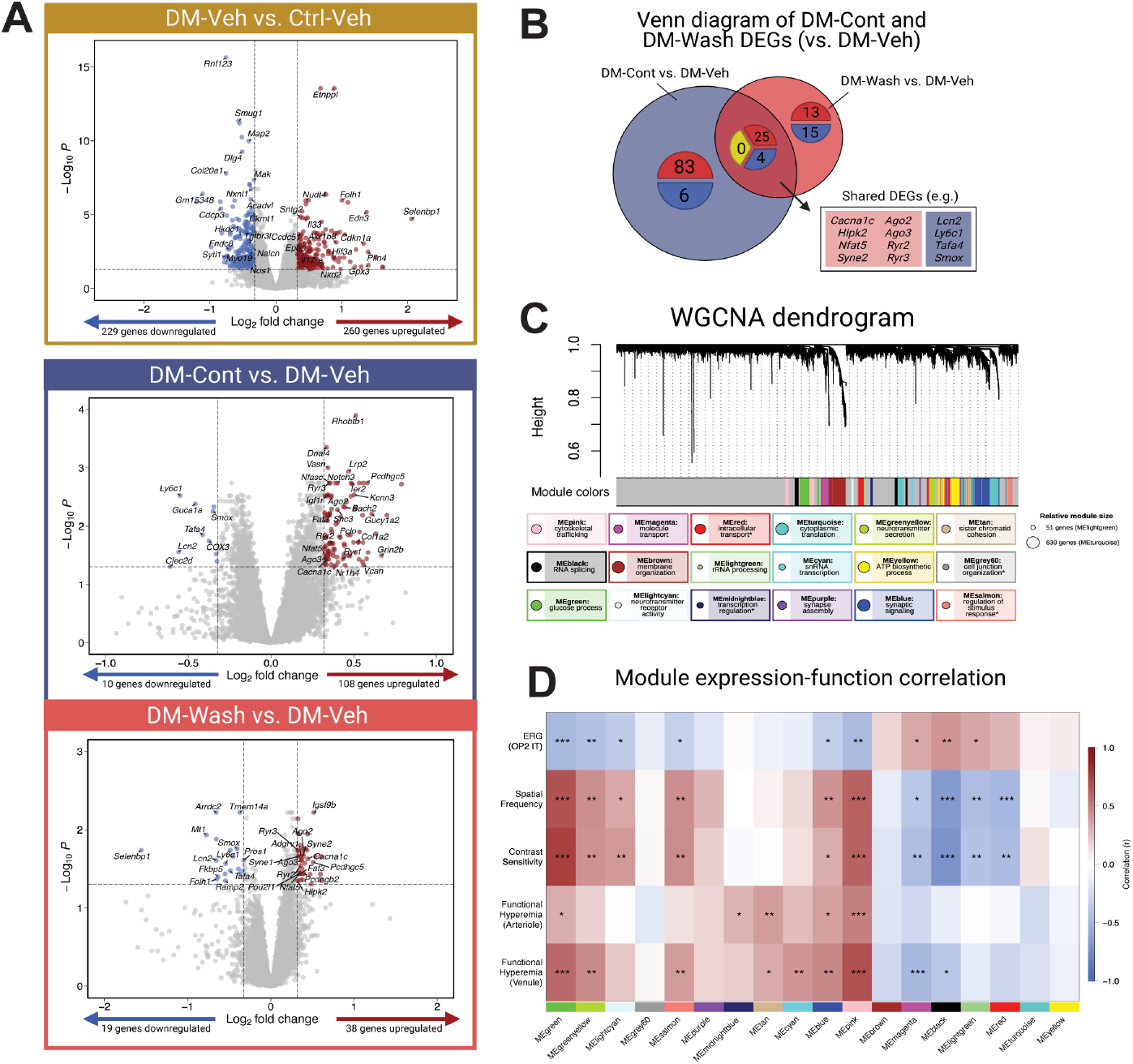
L-DOPA continued and washout treatment resulted in transcriptional changes that correlated with retinal function improvement. (**A**) DM+Veh mice had 260 upregulated and 229 downregulated DEGs compared to Ctrl+Veh mice; |log_2_ fold change| > 0.32, p.adj < 0.05). When comparing DM-Cont to DM-Veh groups, 108 upregulated and 10 downregulated DEGs were identified. When comparing DM-Wash to DM-Veh, 38 upregulated and 19 downregulated genes were highlighted (|log_2_ fold change| > 0.32, p.adj < 0.05). (**B**) Venn diagram of DM-Cont vs. DM-Veh DEGs and DM-Wash vs. DM-Veh DEGs, illustrates shared DEGs including upregulated genes (25) and downregulated genes (4) with no contra-regulated genes (0). (**C**) WGCNA highlighted 18 module eigengenes (MEs) across diabetic and L-DOPA treatment status. Below the dendrogram, each module is summarized in identity (overarching term from GO:BP) and size (relative circle size). Asterisk indicates modules without significant GO:BP terms. (**D**) WGCNA module-function correlation resulted in multiple relevant modules with significant correlations across all functional measures, including MEblue, MEpink, and MEgreen (|r| > 0.30, p < 0.05).

Weighted gene co-expression network analysis (WGCNA) was used to identify modules of co-expressed genes across treatment groups that were correlated with retinal functional outcomes. WGCNA identified 18 modules, or clusters of genes co-expressed across retinal samples (**Fig. 5C**). These modules were then described by top gene ontology (GO) terms related to biological processes (e.g. glucose process, synapse assembly, transcription regulation).

Within each module, the module eigengene (ME) (first principal component of principal component analysis) represents gene expression of co-varying genes per module. When individually correlated with the final week of retinal functional assessments across ERG, OMR, and FH, modules MEgreen, MEblue, and MEpink maintained significant correlation across all functional measures (**Fig. 5D**). Within these modules, only MEblue and MEpink modules were significantly different between DM+Veh and DM+Cont/Wash groups and thus treatment-sensitive (**Fig. 6A, D**). Notably, no modules were significantly different between DM+Cont and DM+Wash.

**Fig. 6.**
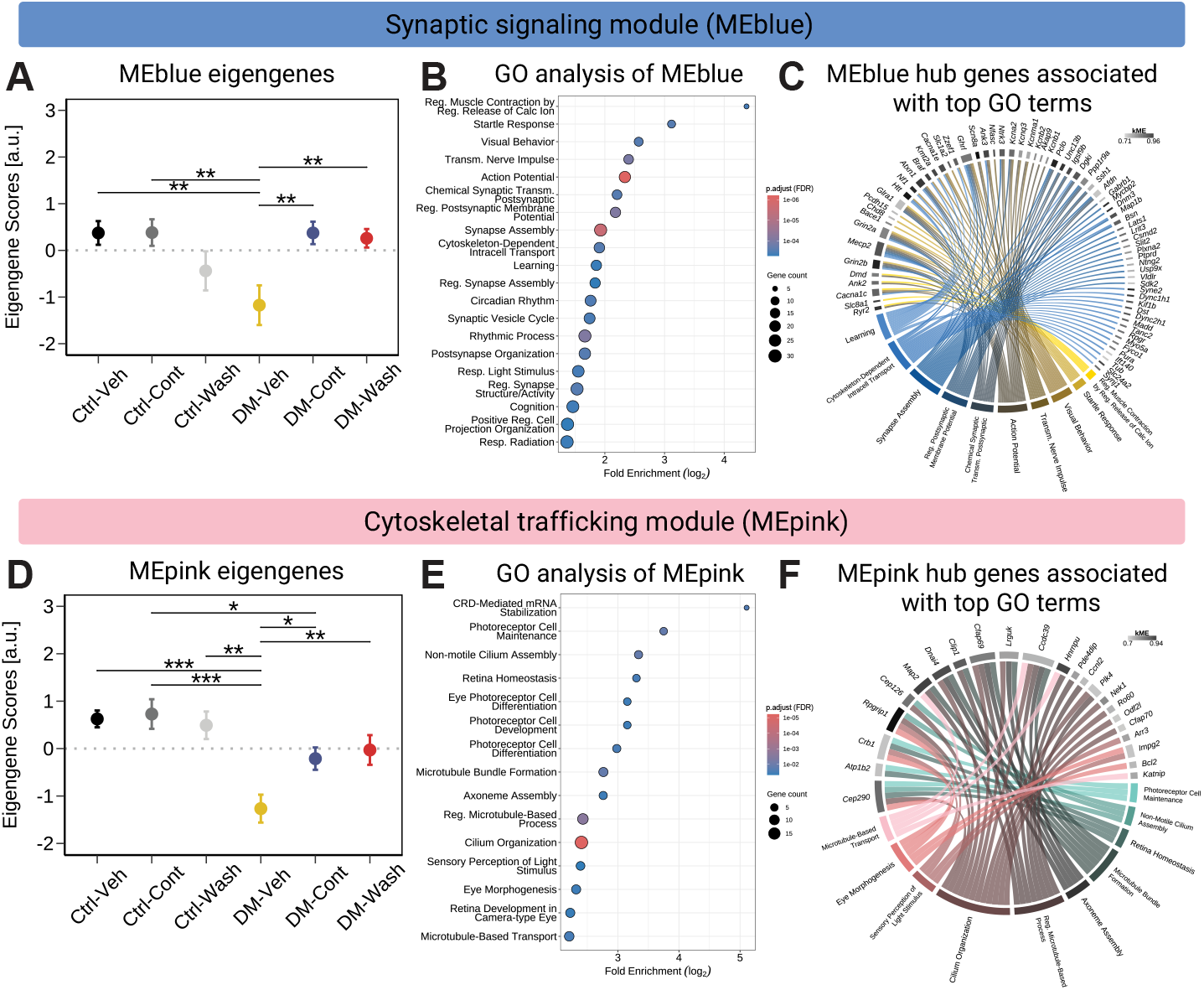
WGCNA modules sensitive to L-DOPA treatment in diabetic mice reflect changes in neuronal signaling, synapses, and cell maintenance. (**A**) Within MEblue, the module eigengene (ME) significantly differed between DM+Veh versus DM+Cont and DM+Wash groups (mean ± SEM, linear model, p.adj < 0.05). (**B**) GO analysis for biological processes of relevant MEblue genes (kME > 0.60) identified signaling and plasticity-related GO terms (FDR-adjusted p < 0.05). (**C**) Chord diagram representing MEblue hub genes (kME > 0.70) shows associations to top 10 enriched GO terms (> 2 gene count). Chord color indicates GO term connection with module-specific gene with kME shown as a separate grayscale bar per gene. (**D**) Similar to MEblue, MEpink significantly differed in DM-Cont and DM-Wash compared to DM-Veh groups (mean ± SEM, linear model, p.adj < 0.05). (**E**) GO analysis for biological processes of relevant MEpink genes (kME > 0.60) identified GO terms related to cytoskeletal dynamics and ciliary function (p.adjust < 0.05). (**F**) Chord diagram representing MEpink hub gene (kME > 0.70) illustrates associations with top 10 enriched GO terms (>2 gene count).

GO analysis of MEblue-specific hubgenes (> 0.7 kME) revealed top terms involving synapse signaling and neurotransmission (e.g. action potential, synapse structure and activity) as well as sensory and rhythmic processes (e.g. response to light stimulus, circadian rhythm) (**Fig. 6A-C**). Interestingly, 20 MEblue hubgenes (> 0.7 kME) (e.g. calcium voltage-gated channel subunit alpha 1 a, *Cacna1c;* ryanodine receptor 2, *Ryr2*) identified as overlapping DEGs shared between DM-Cont vs. Veh and DM-Wash vs. DM-Veh comparisons (**Fig. 6C**). Given the dysregulation of calcium influx in diabetic retinopathy (*31–33*), this shift toward homeostatic expression of calcium channel genes in DM+Cont/Wash suggests L-DOPA-mediated transcriptional stabilization related to calcium signaling. Cell-type percent enrichment analysis of MEblue hubgenes reflected limited (<10%) cell-specific identity, with modest enrichment distributed across neuronal (e.g., bipolar, horizontal, rods, cones) and vascular components (e.g., pericytes and endothelial cells) (**fig. S3, B and D**). Alongside correlation with improvement in diabetic retinal function, this module consists of a broad and sustained change of synaptic and calcium-associated gene expression in the diabetic retina following L-DOPA treatment.

Furthermore, the MEpink module suggested L-DOPA-associated regulation of cytoskeletal components in the diabetic retina. GO analysis of MEpink hubgenes (>0.7 kME) highlighted terms relevant to 1) retinal maintenance (e.g. retina homeostasis, photoreceptor cell maintenance), 2) cilium and axoneme organization (e.g. cilium organization, axoneme assembly), and 3) microtubule-based processes (e.g. microtubule-based transport, microtubule bundle formation) (**Fig. 6D-F**). Notable MEpink hubgenes associated with top GO terms included critical regulators in microtubule stability and dynamics (microtubule-associated protein 2, *Map2;* CAP-Gly domain containing linker protein 1, *Clip1*) (*34–36*), as well as cAMP-signaling and anti-apoptotic survival factors (phosphodiesterase 4D interacting protein, *Pde4dip;* B-cell lymphoma 2, *Bcl2*) (*37, 38*) (**Fig. 6F**). While cell-type percent enrichment analysis of MEpink hubgenes was similarly lacking in a cell-specific identity (<12%), enrichment was primarily neuronal and photoreceptor-driven across rods and cones (f**ig. S3, B and D**). Together, this module supports early cytoskeletal remodeling observed in the diabetic retina (*39, 40*) and suggests a sustained shift to control-like homeostatic expression of cytoskeletal maintenance genes with L-DOPA treatment.

## Discussion

This study demonstrated L-DOPA-mediated mitigation of early functional deficits across the NVU in the diabetic retina and investigated the underlying transcriptional alterations associated with its sustained protective effects. Prior to vascular structural pathology, we show retinal neuronal dysfunction is accompanied by vascular dysfunction in diabetic mice. L-DOPA treatment benefited functional measures of ERG OP IT, spatial frequency, and contrast sensitivity, with protection sustained in diabetic mice for at least two weeks after treatment ended, mirroring clinical work (*10*). Importantly, we demonstrated that L-DOPA also had a direct, and sustained, protective effect on light-induced vascular function in diabetic mice. Analysis of whole retinal transcriptional changes with RNASeq revealed that over half of DEGs in mice with continuous L-DOPA treatment were shared by the L-DOPA washout group. Gene network analysis correlated with L-DOPA-induced improvement of diabetic retinal function, suggesting L-DOPA affects gene networks surrounding synaptic regulation and cytoskeletal maintenance (**Fig. 7**).

**Fig. 7.**
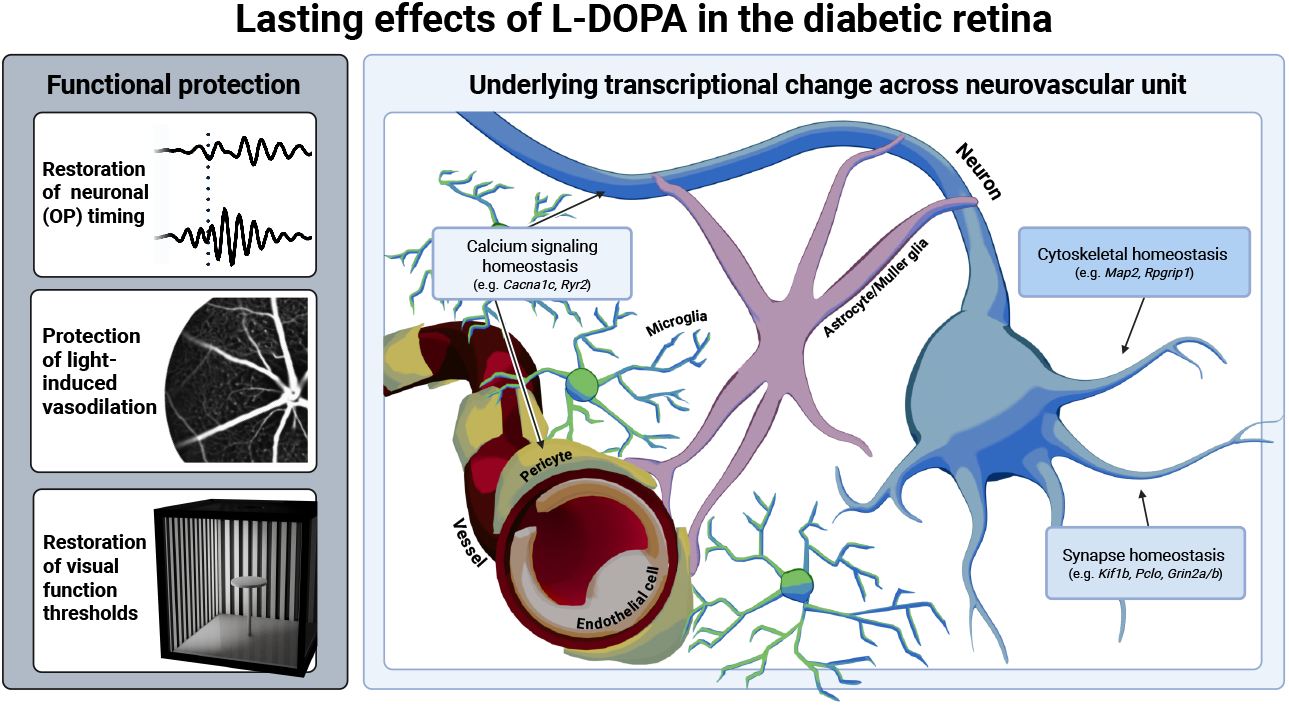
Visual summary of the lasting protective effects of L-DOPA on the diabetic retina across changes in function and associated gene transcription.

### Dopamine benefits neurovascular dysfunction in early DR

While clinical diagnosis has traditionally relied on vascular pathology visible in fundus imaging, there is growing evidence that neuronal deficits precede large-scale vascular change. We have previously focused on neuronal dysfunction as a diagnostic tool for early DR, showing that ERG OP timing, visual acuity, and contrast sensitivity were affected early in diabetic rodents (*8, 9, 11*). In the diabetic retina, we have reported that the dopamine precursor L-DOPA provides functional protection in neuronal dysfunction across diabetic rodent models (*8, 9, 11*) and human participants (*10*). Novel to this study, L-DOPA treatment showed restoration of light-induced retinal vasodilation, supporting the neuroprotective effects of L-DOPA across NVU dysfunction in early DR.

While dopamine is typically described as a retinal neuromodulator, it also acts on vascular and glial components of the NVU. Dopamine receptors from both D1-like (D1R, D5R) and D2-like (D2R, D3R, D4R) families are broadly expressed across most major retinal cell types, including endothelial cells and pericytes. In pericytes, dopamine activation of ATP-sensitive potassium currents serves as a mechanism of metabolic sensing and regulation of vascular tone (*41*). Dopamine has also been associated with anti-angiogenic properties in the retina (*42–44*). Together, these observations support plausible means of L-DOPA-derived modulation of retinal vascular responses.

With protection seen across retinal electrophysiology, visual behavior, and flicker-induced vasodilation, these findings suggest that L-DOPA treatment benefits multiple, interdependent components of the NVU. Reciprocal signaling across the NVU is well-established in retinal pathology (*45–47*). For instance, neuronal *Nrf2* activation in the ischemic retina promotes revascularization through paracrine crosstalk with endothelial cells (*48*). Additionally, elevation of dopamine by L-DOPA is fleeting, reaching its peak within an hour (*49, 50*). Thus, functional benefit seen across the NVU is likely to come from cumulative, interdependent changes across retinal cell types that propagate over time.

### Gene network changes correlate with L-DOPA-induced neuroprotection in DR

While L-DOPA’s relevance spans across neuronal, vascular, and glial components, the biology underlying its protection in the diabetic retina has been unclear. Here, we examined gene network changes associated with L-DOPA-mediated neurovascular protection in the diabetic retina. Using WGCNA analysis, we identified gene modules responsive to L-DOPA treatment in the diabetic retina and correlated with improvements in retinal function. These modules showed heterogenous, cross-NVU identity with limited retinal cell-type enrichment, and notably, none were sensitive to L-DOPA treatment frequency (continued vs. washout). This suggests preservation of protective gene network changes during washout rather than engaging separate, novel changes.

Within the synaptic signaling module (MEblue), L-DOPA treatment was associated with concerted homeostatic expression of signaling and synapse-related genes that correlated with improved retinal function. Prominent among these were genes relevant to intracellular calcium regulation across neurons, glia, and vasculature, including voltage-gated calcium channels and ryanodine receptors (e.g. *Cacna1c, Ryr2*). Dysregulation of calcium signaling is a part of DR’s early pathology, including reduced calcium signaling in presynaptic GABAergic amacrine cells (*31*) as well as hyperglycemia-induced excitotoxic calcium overload (*51, 52*), upregulation of low-voltage T-type calcium channels (*53*), and delayed calcium buffering (*51*). In this context, our findings suggest a nuanced role of L-DOPA on calcium channels in the diabetic retina, potentially restoring balanced calcium influx and cycling to support synaptic function and homeostasis.

Beyond calcium signaling, the synaptic signaling module included other integral genes supporting synapse structure and performance. These included motor proteins that support synaptic function across retinal neurons (e.g. kinesin family member 1b, *Kif1b;* myosin VA, *Myo5a*) (*54, 55*), glutamatergic receptor components crucial to synaptic plasticity and enriched in amacrine and retinal ganglion cells (e.g. glutamate ionotropic receptor NMDA type subunit 2A and 2B, *Grin2a* and *Grin2b*) (*56*), and scaffolding proteins essential for ribbon synapse organization and neurotransmitter release at photoreceptor and bipolar cell synapses (e.g. piccolo presynaptic cytomatrix protein, *Pclo;* bassoon presynaptic cytomatrix protein, *Bsn*) (*57, 58*). Given prior observations of synapse-related gene downregulation in diabetic rodents (*56, 59–61*), coordinated regulation of these synapse-relevant genes suggests that L-DOPA enhances synaptic resilience, with downstream benefit for NVU integrity by stabilizing neuronal activity and reducing stress on glial and vascular components.

L-DOPA sensitivity and correlation with neurovascular functional improvement was also observed in a cytoskeletal trafficking module (MEpink). Core to this module were microtubule-associated genes (e.g. *Map2, Clip1, Pde4dip*), reflecting L-DOPA-induced coordinated regulation of microtubule dynamics, intracellular trafficking, and cellular migration that supports dendritic and synaptic architecture (*34–36, 38*). Notably, *Map2* is known to be reduced in the diabetic retina, indicating early retinal neurodegeneration linked to cytoskeletal dysfunction (*39, 62*). The same module was also enriched for ciliary genes (e.g. RPGR interacting protein 1, *Rpgrip1;* centrosomal protein 290, *Cep290;* Bardet-Biedl syndrome 1, *Bbs1*) essential to photoreceptor maintenance and function (*63– 65*), contributing to the module’s photoreceptor-specific neuronal enrichment.

Additionally, *Bbs1* is required for vascular signaling and mechanotransduction (*66, 67*) and *RPGRIP1* is an emerging potential target within human molecular signatures for proliferative DR (*68*). Collectively, coordinated regulation of microtubule and ciliary gene expression alongside calcium and synaptic signaling genes support a model in which L-DOPA promotes structural and functional plasticity under hyperglycemic stress, strengthening communication across the NVU and contributing to protection of retinal function.

### L-DOPA’s sustained effect in the diabetic retina

In the diabetic retina, we observed improvements in neurovascular function and changes in gene expression after L-DOPA treatment which extended far beyond its expected pharmacokinetic window. L-DOPA’s short half-life is a core limitation to its use as a therapeutic strategy in disorders like Parkinson’s disease (*49*). Even when paired with decarboxylase inhibitors to minimize peripheral metabolism, L-DOPA is found in plasma and cerebrospinal fluid for at most a few hours (*50*). We hypothesize that for L-DOPA to have sustained functional protection weeks beyond treatment, its effect stretches across multiple aspects of the interdependent NVU and may even be independent of dopamine metabolism.

One possible explanation is within the coordinated improvements in neurovascular function, potentially establishing a protective feedback loop across the NVU. As described previously, addressing one or more components of the NVU can encourage stabilization in secondary components. This may, in turn, reinforce preceding benefit and propagate a self-sustaining state rather than transient change. In this way, dopamine’s relevance to all NVU components may encourage a homeostatic feedback loop, producing functional protection that persists beyond transient neurotransmitter-receptor interactions. This hypothesis suggests that the breadth of L-DOPA’s relevance across the NVU not only mediates cross-functional benefit but may also facilitate lasting functional protection.

Separately, recent research on L-DOPA’s role in the melanin synthesis pathway has established 1) L-DOPA is an endogenous ligand to GPR143, a G-protein coupled receptor found in the retinal pigment epithelium (RPE) (*69, 70*), and 2) L-DOPA benefits retinal and visual function in a rodent albinism model lacking functional tyrosinase for synthesizing L-DOPA in the RPE (*71*). While L-DOPA treatment restores reduced retinal dopamine levels in diabetic rodent models (*8, 9*), this research highlights the possibility that the lasting effect of L-DOPA treatment in the diabetic retina could be supported downstream of L-DOPA itself and not exclusively by dopamine. This is similarly postulated for a retrospective study on age-related macular degeneration (AMD) in patients taking L-DOPA, where exogenous L-DOPA significantly delayed AMD development compared to patients without prescription or taking dopaminergic agonists (*72*). However, the role of L-DOPA in AMD remains contested, with other work assessing choroidal neovascularization determining L-DOPA conversion to dopamine as necessary for benefit in *ex vivo* and *in vitro* conditions (*73*). In summary, while the primary metabolic pathways of exogenous L-DOPA in the diabetic retina have yet to be identified, receptors like GPR143 remain a potential player that could contribute to gene expression changes that outlive L-DOPA’s presence in circulation.

This work established sustained neurovascular protection by L-DOPA in the diabetic mouse retina, in line with previous clinical findings (*10*). Here, we have shown L-DOPA-induced protection or restoration across measures of rod-driven inner retinal electrical activity, visual thresholds for spatial frequency and contrast sensitivity, and, importantly, vasodilative response to photic flicker. Using gene expression analysis, we highlighted a potential role of L-DOPA in shaping components of synaptic signaling and cytoskeletal maintenance. Given L-DOPA’s existence as a repurposed drug that provides sustained benefit to neurovascular components vulnerable to hyperglycemia in the retina, it maintains potential as a clinical treatment option to prevent vision-threatening DR. Limitations of this study include its specificity for early stages of DR without proliferation, as well as its short time window that does not assess the long-term effects of continued L-DOPA treatment on the diabetic retina. Future research on L-DOPA’s sustained effect in the diabetic retina should confirm protein-level changes with retinal cell specificity, assess potential sex-specific L-DOPA pharmacokinetics (*74*), and determine L-DOPA’s primary metabolic pathway.

## Materials and Methods

### Animals

Three-month-old male C57BL/6J mice were acquired from Jackson Laboratories (Bar Harbor, ME, USA) and housed in the animal facility at the Joseph Maxwell Cleland Atlanta Veterans Affairs Medical Center (Decatur, GA) under a 12:12-hour (light:dark) cycle with food and water ad libitum. Hyperglycemia was induced via daily i.p. injection of low-dose STZ (50 mg/kg, in citrate buffer, pH 4.5) across five days, with controls receiving equal volumes of citrate buffer vehicle. Blood glucose (BG) was monitored via tail vein blood collection assessed by a handheld glucose meter (Freestyle; Abbott, Abbott Park, IL, USA), with hyperglycemia confirmed by two consecutive BG measurements of >250 mg/dL. BG and body weights were monitored weekly until the end of the study. In the event of successive weight reduction (2% body weight) following hyperglycemia, a single i.p. injection of low-dose insulin (1:10 units of insulin/lactated ringer’s) was administered to prevent weight loss but avoid hypoglycemia. All procedures were approved by the Institutional Animal Care and Use Committee of the Atlanta Veterans Affairs Medical Center and performed in full accordance with the ARVO Statement for the Use of Animals in Ophthalmic and Vision Research.

### Experimental design

Before diabetic induction, baseline measurements of retinal function and visual function were assessed with electroretinography (ERG) and optomotor response (OMR), respectively (**Figure 1**). Starting four weeks post-hyperglycemia, ERGs and OMRs were collected weekly to identify initial retinal and visual functional deficits. Following detection of functional deficits, mice were randomized to daily oral treatment of L-DOPA or vehicle for two weeks. L-DOPA-treated mice were then further divided into two groups: continued L-DOPA treatment (Cont) or vehicle only to create a L-DOPA washout phase (Wash). Group allocation was the following: Control + Vehicle (Ctrl+Veh, n=9), Control + Continued L-DOPA (Ctrl+Cont, n=9), Control + Washout L-DOPA (Ctrl+Wash, n=6), Diabetic + Vehicle (DM+Veh, n=12-15), Diabetic + Continued L-DOPA (DM+Cont n=12-15), Diabetic + Washout L-DOPA (DM+Wash, n=7-11). ERGs and OMRs were collected weekly during the following two weeks of ‘washout’/’continued’ L-DOPA treatment. In the final week, functional hyperemia measurements were acquired, followed by euthanasia via cervical dislocation. Retinas from left eyes were flash-frozen for RNA isolation while retinas from right eyes were PFA-fixed for future immunohistochemistry.

### Oral L-DOPA treatment

L-DOPA (Sigma-Aldrich, St. Louis, MO) was suspended in sweetened condensed milk (California Farms, Santini Foods, Inc.) and fed to mice under dim-light conditions in individual static cages. As previously described (*75*), L-DOPA treatment was prepared fresh daily under dim-light conditions with L-DOPA (20 mg/kg at 20 mg/mL) and carbidopa monohydrate (5 mg/kg at 5 mg/mL), the latter maximizing delivery of L-DOPA through prevention of premature peripheral L-DOPA oxidation (*49, 76*). Once combined with sweetened condensed milk, the treatment was thoroughly mixed until homogenous and immediately distributed via petri dish. Consumption of oral L-DOPA typically occurred under a minute and was confirmed by visual inspection. Oral L-DOPA treatment was administered after functional assessments on testing days. One cohort was treated with L-DOPA/carbidopa at one-quarter of the original dosage, showed no functional differences in continuously treated groups, and was combined with the other groups.

### Electroretinograms (ERGs)

For retinal function assessment, mice were dark-adapted overnight and prepared for ERG recordings under dim red light, as previously described (*8, 9, 11*). Following anesthesia (ketamine [80 mg/kg] and xylazine [16 mg/kg]), pupils were dilated (1% tropicamide), corneal surfaces were anesthetized (0.5% tetracaine HCl), and body temperature was maintained with a heating pad. Custom-made gold loops placed at the corneal surface served as the active electrode, while subdermal cheek and tail electrodes operated as reference and ground electrodes, respectively. Throughout recordings, eye drops (Systane Ultra; Alcon Laboratories, Inc., Fort Worth, TX) maintained corneal hydration and optimal electrode-cornea contact. Electrical responses to full-field flash stimuli presented within a Ganzfeld dome were recorded and amplified with a signal-averaging ERG system (UTAS Bigshot; LKC Technologies, Gaithersburg, MD). Using a custom five-step flash protocol, white light stimuli within a dark-adapted series were presented (-3.0, -1.7, -0.6, 1.5, 1.9 log cd s/m^2^). After ERG acquisition, mice were administered atipamezole hydrochloride (1 mg/kg; Antisedan; Pfizer Animal Health, New York, NY) to facilitate anesthesia recovery. Oscillatory potentials were filtered digitally with a high-pass filter (75-350 Hz) using custom MATLAB software (*77*). Data from only one eye (the eye with the largest oscillatory potential amplitude in response to -3.0 log cd s/m2 flash) was analyzed at each flash intensity per animal. Amplitudes and implicit times of ERG a-waves (photoreceptor function), b-waves (bipolar cell function), and oscillatory potentials (amacrine function) were measured, as previously published (*9–11*).

### Optomotor response (OMRs)

Visual function of mice was assessed with a virtual optomotor system (OptoMotry system; Cerebral-Mechanics, Lethbridge, AB, Canada). Mice were placed on the central pedestal of a virtual reality chamber consisting of four surrounding computer monitors, as previously described (*8, 9*). Mice were given one minute of environmental acclimation before monitors were turned on to display vertical sine wave gratings rotating at 12 deg/s speed. A staircase paradigm separately probed spatial frequency (SF)(100% contrast) and contrast sensitivity (CS)(0.103 cycles/degree) thresholds. With active monitoring by the experimenter, reflexive head movements (tracking) and mouse head alignment with the center of the virtual cylinder was maintained.

### Functional hyperemia (FH)

To assess retinal vascular function, confocal scanning laser ophthalmoscopy (Heidelberg Spectralis; HRA+OCT; Heidelberg Engineer, Carlsbad, CA) was performed during photopic stimulation. Following anesthesia and eye preparation as described for ERGs, retinal blood vessels were visualized via intraperitoneal injection of indocyanine green dye (ICG, 18.75 mg/kg; IC-Green; Akorn, Lake Forest, IL). Square wave 12 Hz green light (480-600 nm) was presented through a fiber optic bundle and reflected off a 45-degree angle prism mirror (TS Cold Mirror; Edmund Optics, Barrington, NJ) to stimulate the retina without interrupting image acquisition. Light was presented in light-adapted conditions, with luminance at 4000-5000 lux at eye surface. ICG angiography video was acquired at high resolution with a widefield 55-degree scan angle. A functional hyperemia recording consisted of a 10 second pre-flicker baseline, 15 seconds of flicker stimulation, and 10 seconds of post-flicker. In the event of a suboptimal trial (ex: movement), another trial was acquired following two minutes of separation.

To quantify flicker-induced vasodilation, vessel caliber kymographs were generated from ICG fundus video sequences (ImageJ, U. S. National Institutes of Health, Bethesda, Maryland, USA) and aligned within-stack before regions of interest representing optic disc size were generated. Using first-order arteriole or venule, single cross-sections were measured one optic disc distance away from the optic nerve. After thresholding, kymographs were generated from two randomly selected arterioles and venules per eye. Vessel identity relied on morphology, depth, and alternating pattern (*78, 79*). To determine percent vasodilation, vessel diameters at baseline (0-10 seconds) and stimulation (12.5-22.5 seconds) were averaged in a custom MATLAB script and compared following a moving window average smoothing algorithm (Mathworks; Natick, MA, US). Reported percentages were determined by averaging vasodilation values for arterioles and venules.

### RNA sequencing and data preparation

For RNASeq bulk analysis, retinas from left eyes were collected in ambient lighting between 10:00 AM and 2:00 PM, flash-frozen on dry ice, and stored at -80 C. Mice on continuous L-DOPA treatment were administered L-DOPA 15 minutes prior to enucleation. Total RNA was extracted (RNeasy, QIAGEN; Catalog no. 74106) following the manufacturer’s protocol and sent to Admera Health (South Plainfield, NJ) for bulk sequencing and alignment. Following quality control by thresholding for RNA Integrity Number (RIN>7), mRNA was prepared with NEB Next Ultra (II) Directional Kit with Poly A Selection and sequenced (2x150 bp) using the Illumina platform at 20 million reads/direction. Reads were aligned with STAR Aligner (v2.7.1a) and gene counts/sample were determined with HTseq-count (v 0.11.2).

### Differential gene expression analysis

Transcripts with fewer than 40 counts were filtered out, leaving 13,665 total transcripts for downstream analysis. Normalization of raw read counts and differential expression analysis was performed with *DESeq2* R package (v1.49.2). Counts were normalized with the median-of-ratios method, and a design formula specifying condition was used to fit a negative binomial generalized linear model for each gene. To address variability due to batch effects observed in bulk tissue RNA-seq data, the *RemoveBatchEffect* function from the *limma* package in R was applied for correction. Principle component analysis (PCA) of normalized, batch-corrected data was used for outlier identification by calculating the Mahalanobis distance of each point, with samples falling outside of 95% confidence ellipse in PCA space removed in two iterative rounds and PCA re-evaluated after each round. In total, seven samples were detected as outliers and removed from the study.

Differentially expressed genes (DEGs) were determined using cut-offs for Benjamini-Hochberg adjusted p-value (<0.05) and absolute log_2_ fold change (>0.32). R package *EnhancedVolcano* (v1.27.0) was used for DEG volcano visualization.

### Weighted gene co-expression network analysis

Weighted gene co-expression network analysis (WGCNA) was performed using the *WGCNA* R package (v1.73). Raw read counts were first normalized with *DESeq2* (v1.49.2) via median-of-ratios and a design formula of condition and batch. Normalized gene counts were transformed to stabilize variance and residual batch effects were removed using the *limma* R package (v3.65.1). Prior to network construction, lowly expressed genes were filtered out to reduce noise. In constructing a weighted co-expression network with WGCNA, signed adjacency was used with *biweight midcorrelation*, and a soft-thresholding power of 8 was selected to achieve scale-free topology fit index (R2>0.8). Modules of co-expressed genes were identified using hierarchical clustering with a minimum module size of 50, a maximum module size of 13,665, a deep split size of 4, a merge cut height of 0.25, and mean-based topological overlap matrix. From the network, module eigengenes (MEs) per sample were correlated with raw functional data using Pearson correlation to assess the relationship between module-specific co-expression patterns and retinal function. Cut-offs for correlation strength (|r|>0.3) and statistical significance (unadjusted p<0.05) were used to identify relevant module-trait associations.

### Gene ontology analysis

Gene ontology (GO) enrichment analysis for biological processes was performed for thresholded genes within WGCNA modules (>0.6 kME) using *clusterProfiler* R package (v4.17.0) and *enricher* function. Gene sets were obtained from the MSigDB database with the *msigdbr* R package, filtered for Mus Musculus species and the GO:BP (M5) subcollection. Benjamini-Hochberg false discovery rate (FDR) adjustment to p-values was used (<0.05). R package *circlize* (v0.4.16) was used for chord diagram visualization.

### Retinal cell-type percent enrichment analysis

To evaluate cell-type specificity of WGCNA modules, we performed enrichment analysis across NVU class and major retinal cell type. Using the single cell RNAseq dataset from the Mouse Retina Cell Atlas (*80*), unique gene sets were formed across broad NVU classes (e.g. neuronal, glial, microglial, and vascular) and major cell type classes (e.g. bipolar, amacrine, ganglion, Muller glia, astrocytes, microglia, endothelial cells, pericytes, rods, and cones). Genes were filtered for significant differential expression (log2 fold change > 0.25, adjusted p < 0.05, top 1000 genes per major cell type; top 2000 genes per NVU class), and only retained if unique within class. For each WGCNA-derived module, percent enrichment was determined by dividing the number of cell type-specific or NVU class-specific genes by the total number of thresholded hubgenes (>0.7 kME) present in each module.

### Functional data statistical analysis

Analysis of functional data was performed using statistical software (GraphPad Prism v10.6.1, GraphPad Software, San Diego, CA). Longitudinal ERG and OMR datasets were analyzed using a two-way mixed effects model, while functional hyperemia measures were analyzed with a one-way analysis of variance (ANOVA). All functional measure statistics reported are interaction effects followed by Tukey’s multiple comparison test.

Statistical significance threshold is indicated by *p < 0.05, **p < 0.01, **p < 0.01, ***p < 0.001, ****p < 0.0001. Significance symbols used in longitudinal data reflects group comparisons: (*) DM+Veh vs. Ctrl+Veh, (†) DM+Veh vs. DM+Cont, (‡) DM+Veh vs. DM+Wash, (§) Ctrl+Veh vs. DM+Cont, (#) Ctrl+Veh vs. DM+Wash. Figures show mean ± SEM for each group.

## Acknowledgments

Fig. 1, 2, 4, 5, and 6 included assets from BioRender.

## Funding

Department of Veterans Affairs Merit Award RX002615, RX003825 (MTP)

Department of Veterans Affairs Research Career Scientist Award RX003134 (MTP)

Department of Veterans Affairs Career Development Award BX005304 (KLB)

NIH T32EY007092 (EC)

NEI Core Grant P30EY006360

Research to Prevent Blindness Award to Department of Ophthalmology, Emory University

The George W. Woodruff School of Mechanical Engineering Faculty Fellowship at Georgia Tech (LBW)

## Author contributions

Conceptualization: EC, MTP

Methodology: EC, MTP

Software: EC, CL, LBW

Investigation: EC, KLB, CL

Visualization: EC

Supervision: MTP, LBW

Writing—original draft: EC

Writing—review & editing: EC, CL, KLB, LBW, MTP

## Data and materials availability

The dataset generated and analyzed during this study will be publicly available at the National Center for Biotechnology Information Gene Expression Omnibus after peer-reviewed publication.

## Supplementary Materials

Figs. S1 to S3

Data S1 to S2

## References

1. R. Simó, A. W. Stitt, T. W. Gardner, Neurodegeneration in diabetic retinopathy: does it really matter? Diabetologia 61, 1902–1912 (2018).

2. L. Ji, H. Tian, K. A. Webster, W. Li, Neurovascular regulation in diabetic retinopathy and emerging therapies. Cell Mol Life Sci 78, 5977–5985 (2021).

3. S. R. Levine, P. Sapieha, S. Dutta, J. K. Sun, T. W. Gardner, It is time for a moonshot to find “Cures” for diabetic retinal disease. Progress in Retinal and Eye Research 90, 101051 (2022).

4. E. A. Lundeen, Z. Burke-Conte, D. B. Rein, J. S. Wittenborn, J. Saaddine, A. Y. Lee, A. D. Flaxman, Prevalence of Diabetic Retinopathy in the US in 2021. JAMA Ophthalmology 141, 747–754 (2023).

5. A. N. Kollias, M. W. Ulbig, Diabetic retinopathy: Early diagnosis and effective treatment. Dtsch Arztebl Int 107, 75–83; quiz 84 (2010).

6. L. Z. Heng, O. Comyn, T. Peto, C. Tadros, E. Ng, S. Sivaprasad, P. G. Hykin, Diabetic retinopathy: pathogenesis, clinical grading, management and future developments. Diabet Med 30, 640–650 (2013).

7. H. A. Hancock, T. W. Kraft, Oscillatory potential analysis and ERGs of normal and diabetic rats. Invest Ophthalmol Vis Sci 45, 1002–1008 (2004).

8. M. H. Aung, H. N. Park, M. K. Han, T. S. Obertone, J. Abey, F. Aseem, P. M. Thule, P. M. Iuvone, M. T. Pardue, Dopamine deficiency contributes to early visual dysfunction in a rodent model of type 1 diabetes. J Neurosci 34, 726–736 (2014).

9. K. Chesler, C. Motz, H. Vo, A. Douglass, R. S. Allen, A. J. Feola, M. T. Pardue, Initiation of L-DOPA Treatment After Detection of Diabetes-Induced Retinal Dysfunction Reverses Retinopathy and Provides Neuroprotection in Rats. Transl Vis Sci Technol 10, 8 (2021).

10. C. T. Motz, K. C. Chesler, R. S. Allen, K. L. Bales, L. M. Mees, A. J. Feola, A. Y. Maa, D.E. Olson, P. M. Thule, P. M. Iuvone, A. M. Hendrick, M. T. Pardue, Novel Detection and Restorative Levodopa Treatment for Preclinical Diabetic Retinopathy. Diabetes 69, 1518– 1527 (2020).

11. M. K. Kim, M. H. Aung, L. Mees, D. E. Olson, N. Pozdeyev, P. M. Iuvone, P. M. Thule, M. T. Pardue, Dopamine Deficiency Mediates Early Rod-Driven Inner Retinal Dysfunction in Diabetic Mice. Investigative Ophthalmology & Visual Science 59, 572–581 (2018).

12. H. Albertos-Arranz, N. Martínez-Gil, X. Sánchez-Sáez, J.C. Molina-Martín, P. Lax, N. Cuenca, Neuronal Degeneration and Glial Activation in the Absence of Vascular Changes in Human Retinas of Patients With Diabetes. Invest Ophthalmol Vis Sci 66, 53 (2025).

13. C. J. Pournaras, E. Rungger-Brändle, C. E. Riva, S. H. Hardarson, E. Stefansson, Regulation of retinal blood flow in health and disease. Prog Retin Eye Res 27, 284–330 (2008).

14. J. R. Hombrebueno, M. Chen, R. G. Penalva, H. Xu, Loss of Synaptic Connectivity, Particularly in Second Order Neurons Is a Key Feature of Diabetic Retinal Neuropathy in the Ins2Akita Mouse. PLOS ONE 9, e97970 (2014).

15. A. Mandecka, J. Dawczynski, M. Blum, N. Müller, C. Kloos, G. Wolf, W. Vilser, H. Hoyer, U. A. Müller, Influence of Flickering Light on the Retinal Vessels in Diabetic Patients. Diabetes Care 30, 3048–3052 (2007).

16. G. Garhöfer, C. Zawinka, H. Resch, P. Kothy, L. Schmetterer, G. T. Dorner, Reduced response of retinal vessel diameters to flicker stimulation in patients with diabetes. Br J Ophthalmol 88, 887–891 (2004).

17. A. R. Nippert, E. A. Newman, Regulation of blood flow in diabetic retinopathy. Vis Neurosci 37, E004 (2020).

18. A. I. Jobling, U. Greferath, M. A. Dixon, P. Quiriconi, B. Eyar, A. K. van Koeverden, S. A. Mills, K. A. Vessey, B. V. Bui, E. L. Fletcher, Microglial regulation of the retinal vasculature in health and during the pathology associated with diabetes. Progress in Retinal and Eye Research 106, 101349 (2025).

19. M. G. Rossino, M. Dal Monte, G. Casini, Relationships Between Neurodegeneration and Vascular Damage in Diabetic Retinopathy. Front Neurosci 13, 1172 (2019).

20. N. Nishikiori, M. Osanai, H. Chiba, T. Kojima, Y. Mitamura, H. Ohguro, N. Sawada, Glial cell-derived cytokines attenuate the breakdown of vascular integrity in diabetic retinopathy. Diabetes 56, 1333–1340 (2007).

21. C. Meng, C. Gu, S. He, T. Su, T. Lhamo, D. Draga, Q. Qiu, Pyroptosis in the Retinal Neurovascular Unit: New Insights Into Diabetic Retinopathy. Front. Immunol. 12 (2021).

22. S. Roy, G. D. Field, Dopaminergic modulation of retinal processing from starlight to sunlight. Journal of Pharmacological Sciences 140, 86–93 (2019).

23. C. R. Jackson, G.-X. Ruan, F. Aseem, J. Abey, K. Gamble, G. Stanwood, R. D. Palmiter, P.M. Iuvone, D. G. McMahon, Retinal Dopamine Mediates Multiple Dimensions of Light-Adapted Vision. J Neurosci 32, 9359–9368 (2012).

24. M. Goel, S. C. Mangel, Dopamine-Mediated Circadian and Light/Dark-Adaptive Modulation of Chemical and Electrical Synapses in the Outer Retina. Front. Cell. Neurosci. 15 (2021).

25. C. Nishimura, K. Kuriyama, Alterations in the retinal dopaminergic neuronal system in rats with streptozotocin-induced diabetes. J Neurochem 45, 448–455 (1985).

26. G. T. Dorner, G. Garhöfer, K. H. Huemer, C. E. Riva, M. Wolzt, L. Schmetterer, Hyperglycemia affects flicker-induced vasodilation in the retina of healthy subjects. Vision Research 43, 1495–1500 (2003).

27. A. Mishra, E. A. Newman, Aminoguanidine reverses the loss of functional hyperemia in a rat model of diabetic retinopathy. Front Neuroenergetics 3 (2011).

28. H. B. Haupt, J. M. Nickerson, J. H. Boatright, M. T. Pardue, K. L. Bales, Effects of Treadmill Exercise on Retinal Vascular Morphology, Function, and Circulating Immune Factors in a Mouse Model of Retinal Degeneration. Invest Ophthalmol Vis Sci 66, 65 (2025).

29. X. Deng, K. Liu, T. Zhu, D. Guo, X. Yin, L. Yao, Z. Ding, J. Ye, P. Li, Dynamic inverse SNR-decorrelation OCT angiography with GPU acceleration. Biomed Opt Express 13, 3615–3628 (2022).

30. T. Son, G. Ma, X. Yao, Functional OCT reveals anisotropic changes of retinal flicker-evoked vasodilation. Opt Lett 49, 2121–2124 (2024).

31. J. M. Moore-Dotson, E. D. Eggers, Reductions in Calcium Signaling Limit Inhibition to Diabetic Retinal Rod Bipolar Cells. Invest. Ophthalmol. Vis. Sci. 60, 4063–4073 (2019).

32. A. Saadane, Y. Du, W. B. Thoreson, M. Miyagi, E. M. Lessieur, J. Kiser, X. Wen, B. A. Berkowitz, T. S. Kern, Photoreceptor Cell Calcium Dysregulation and Calpain Activation Promote Pathogenic Photoreceptor Oxidative Stress and Inflammation in Prodromal Diabetic Retinopathy. Am J Pathol 191, 1805–1821 (2021).

33. S. Z. Haider, N. P. Sadanandan, P. G. Joshi, B. Mehta, Early Diabetes Induces Changes in Mitochondrial Physiology of Inner Retinal Neurons. Neuroscience 406, 140–149 (2019).

34. I. Barbiero, D. Peroni, P. Siniscalchi, L. Rusconi, M. Tramarin, R. De Rosa, P. Motta, M. Bianchi, C. Kilstrup-Nielsen, Pregnenolone and pregnenolone-methyl-ether rescue neuronal defects caused by dysfunctional CLIP170 in a neuronal model of CDKL5 Deficiency Disorder. Neuropharmacology 164, 107897 (2020).

35. F. Larti, K. Kahrizi, L. Musante, H. Hu, E. Papari, Z. Fattahi, N. Bazazzadegan, Z. Liu, M. Banan, M. Garshasbi, T. F. Wienker, H. H. Ropers, N. Galjart, H. Najmabadi, A defect in the CLIP1 gene (CLIP-170) can cause autosomal recessive intellectual disability. Eur J Hum Genet 23, 331–336 (2015).

36. R. A. DeGiosio, M. J. Grubisha, M. L. MacDonald, B. C. McKinney, C. J. Camacho, R. A. Sweet, More than a marker: potential pathogenic functions of MAP2. Front Mol Neurosci 15, 974890 (2022).

37. M. Vogler, Y. Braun, V. M. Smith, M.-A. Westhoff, R. S. Pereira, N. M. Pieper, M. Anders, M. Callens, T. Vervliet, M. Abbas, S. Macip, R. Schmid, G. Bultynck, M. J. Dyer, The BCL2 family: from apoptosis mechanisms to new advances in targeted therapy. Sig Transduct Target Ther 10, 91 (2025).

38. W. Gu, H. Li, W. Yuan, X. Fu, R. Wang, X. Xu, X. Liao, L. Liu, B. Pan, J. Tian, H. Yuan, Y. Huang, T. Lu, Dysfunction of PDE4DIP contributes to LVNC development by regulating cell polarity, skeleton, and energy metabolism via Rho-ROCK pathway. Genes & Diseases 12, 101568 (2025).

39. M. E. Marin Castano, M. Ruggeri, D. DeBuc, F. Penha, X.-R. Huang, Retina, Modification in the retinal nerve fiber layer thickness and microtubule-associated proteins (MAPs) expression in diabetic mice. Investigative Ophthalmology & Visual Science 55, 4910 (2014).

40. A. T. Patrick, W. He, J. Madu, S. R. Sripathi, S. Choi, K. Lee, F. P. Samson, F. L. Powell, M. Bartoli, D. Jee, D. R. Gutsaeva, W. J. Jahng, Mechanistic dissection of diabetic retinopathy using the protein-metabolite interactome. J Diabetes Metab Disord 19, 829–848 (2020).

41. D. M. Wu, H. Kawamura, Q. Li, D. G. Puro, Dopamine activates ATP-sensitive K+ currents in rat retinal pericytes. Visual Neuroscience 18, 935–940 (2001).

42. R. S. Allen, C. T. Khayat, A. J. Feola, A. S. Win, A. R. Grubman, K. C. Chesler, L. He, J. A. Dixon, T. S. Kern, P. M. Iuvone, P. M. Thule, M. T. Pardue, Diabetic rats with high levels of endogenous dopamine do not show retinal vascular pathology. Front Neurosci 17, 1125784 (2023).

43. I. Veernala, A. S. Cuamatzi-Castelan, A. Rajesh, J. Gong, G. J. D’Souza, J. A. Lavine, Levodopa Suppresses Choroidal Neovascularization Through a Tyrosinase-Dependent Dual Mechanism. Invest Ophthalmol Vis Sci 67, 8 (2026).

44. S. Upreti, S. Sen, T. C. Nag, M. P. Ghosh, Insulin like growth factor-1 works synergistically with dopamine to attenuate diabetic retinopathy by downregulating vascular endothelial growth factor. Biomed Pharmacother 149, 112868 (2022).

45. W. Xu, L.-J. Cui, X.-Y. Yang, X.-Y. Cui, J. Guo, G.-X. Xu, Impaired pericyte-Müller glia interaction via PDGFRβ suppression aggravates photoreceptor loss in a rodent model of light-induced retinal injury. Int J Ophthalmol 17, 1800–1808 (2024).

46. S. A. Mills, A. I. Jobling, M. A. Dixon, B. V. Bui, K. A. Vessey, J. A. Phipps, U. Greferath, G. Venables, V. H. Y. Wong, C. H. Y. Wong, Z. He, F. Hui, J. C. Young, J. Tonc, E. Ivanova, B. T. Sagdullaev, E. L. Fletcher, Fractalkine-induced microglial vasoregulation occurs within the retina and is altered early in diabetic retinopathy. Proc Natl Acad Sci U S A 118, e2112561118 (2021).

47. D. Rodriguez, K. A. Church, A. N. Pietramale, S. M. Cardona, D. Vanegas, C. Rorex, M. C. Leary, I. A. Muzzio, K. R. Nash, A. E. Cardona, Fractalkine isoforms differentially regulat amicroglia-mediated inflammation and enhance visual function in the diabetic retina. J Neuroinflammation 21, 42 (2024).

48. Y. Wei, J. Gong, Z. Xu, R. K. Thimmulappa, K. L. Mitchell, D. S. Welsbie, S. Biswal, E. J. Duh, Nrf2 in ischemic neurons promotes retinal vascular regeneration through regulation of semaphorin 6A. Proceedings of the National Academy of Sciences 112, E6927–E6936 (2015).

49. J. G. Nutt, Pharmacokinetics and pharmacodynamics of levodopa. Mov Disord 23 Suppl 3, S580–584 (2008).

50. J. P. Hammerstad, W. R. Woodward, P. Gliessman, B. Boucher, J. G. Nutt, L-dopa pharmacokinetics in plasma and cisternal and lumbar cerebrospinal fluid of monkeys. Ann Neurol 27, 495–499 (1990).

51. Y.-C. Wang, L. Wang, Y.-Q. Shao, S.-J. Weng, X.-L. Yang, Y.-M. Zhong, Exendin-4 promotes retinal ganglion cell survival and function by inhibiting calcium channels in experimental diabetes. iScience 26 (2023).

52. E. A. Pereira da Silva, M. Martín-Aragón Baudel, J. Hong, P. Bartels, M. F. Navedo, M. Nieves-Cintrón, “Vascular CaV1.2 channels in diabetes” in Current Topics in Membranes, M. Sturek, Ed. (Academic Press, 2022; https://www.sciencedirect.com/science/article/pii/S1063582322000126)vol. 90 of Ion Transport and Membrane Interactions in Vascular Health and Disease, pp. 65–93.

53. D. E. Duzhyy, V. Y. Viatchenko-Karpinski, E. V. Khomula, N. V. Voitenko, P. V. Belan, Upregulation of T-Type Ca2+ Channels in Long-Term Diabetes Determines Increased Excitability of a Specific Type of Capsaicin-Insensitive DRG Neurons. Mol Pain 11, s12990-015-0028-z (2015).

54. C. Desnos, S. Huet, F. Darchen, “Should I stay or should I go?”: myosin V function in organelle trafficking. Biol Cell 99, 411–423 (2007).

55. S. Chen, M. Han, W. Chen, Y. He, B. Huang, P. Zhao, Q. Huang, L. Gao, X. Qu, X. Li, KIF1B promotes glioma migration and invasion via cell surface localization of MT1-MMP. Oncol Rep 35, 971–977 (2016).

56. J. C. M. Lau, R. A. Kroes, J. R. Moskal, R. A. Linsenmeier, Diabetes changes expression of genes related to glutamate neurotransmission and transport in the Long-Evans rat retina. Mol Vis 19, 1538–1553 (2013).

57. D. Ivanova, A. Dirks, A. Fejtova, Bassoon and piccolo regulate ubiquitination and link presynaptic molecular dynamics with activity -regulated gene expression. J Physiol 594, 5441–5448 (2016).

58. R. A. Voorn, C. Vogl, Molecular Assembly and Structural Plasticity of Sensory Ribbon Synapses-A Presynaptic Perspective. Int J Mol Sci 21, 8758 (2020).

59. F. I. Baptista, M. J. Pinto, F. Elvas, T. Martins, R. D. Almeida, A. F. Ambrósio, Diabetes induces changes in KIF1A, KIF5B and dynein distribution in the rat retina: Implications for axonal transport. Experimental Eye Research 127, 91–103 (2014).

60. H. Ramos, P. Bogdanov, R. Simó, A. Deàs-Just, C. Hernández, Transcriptomic Analysis Reveals That Retinal Neuromodulation Is a Relevant Mechanism in the Neuroprotective Effect of Sitagliptin in an Experimental Model of Diabetic Retinopathy. International Journal of Molecular Sciences 24 (2022).

61. J. M. Gaspar, F. I. Baptista, J. Galvão, A. F. Castilho, R. A. Cunha, A. F. Ambrósio, Diabetes differentially affects the content of exocytotic proteins in hippocampal and retinal nerve terminals. Neuroscience 169, 1589–1600 (2010).

62. H. Zhu, W. Zhang, Y. Zhao, X. Shu, W. Wang, D. Wang, Y. Yang, Z. He, X. Wang, Y. Ying, GSK3β-mediated tau hyperphosphorylation triggers diabetic retinal neurodegeneration by disrupting synaptic and mitochondrial functions. Mol Neurodegener 13, 62 (2018).

63. R. A. Rachel, T. Li, A. Swaroop, Photoreceptor sensory cilia and ciliopathies: focus on CEP290, RPGR and their interacting proteins. Cilia 1, 22 (2012).

64. K. N. Rao, W. Zhang, L. Li, C. Ronquillo, W. Baehr, H. Khanna, Ciliopathy-associated protein CEP290 modifies the severity of retinal degeneration due to loss of RPGR. Hum Mol Genet 25, 2005–2012 (2016).

65. R. K. Raghupathy, X. Zhang, F. Liu, R. H. Alhasani, L. Biswas, S. Akhtar, L. Pan, C. B. Moens, W. Li, M. Liu, B. N. Kennedy, X. Shu, Rpgrip1 is required for rod outer segment development and ciliary protein trafficking in zebrafish. Sci Rep 7, 16881 (2017).

66. J. Jiang, J. J. Reho, S. Bhattarai, I. Cherascu, A. Hedberg-Buenz, K. J. Meyer, F. Tayyari, A.J. Rauckhorst, D. F. Guo, D. A. Morgan, E. B. Taylor, M. G. Anderson, A. V. Drack, K. Rahmouni, Endothelial BBSome is essential for vascular, metabolic, and retinal functions. Mol Metab 53, 101308 (2021).

67. J. J. Reho, D.-F. Guo, D. A. Morgan, K. Rahmouni, Smooth Muscle Cell-Specific Disruption of the BBSome Causes Vascular Dysfunction. Hypertension 74, 817–825 (2019).

68. D. Shao, S. He, Z. Ye, X. Zhu, W. Sun, W. Fu, T. Ma, Z. Li, Identification of potential molecular targets associated with proliferative diabetic retinopathy. BMC Ophthalmol 20, 143 (2020).

69. B. S. McKay, Pigmentation and Vision: Is GPR143 in Control? J Neurosci Res 97, 77–87 (2019).

70. Y. Goshima, D. Masukawa, Y. Kasahara, T. Hashimoto, A. C. Aladeokin, l-DOPA and Its Receptor GPR143: Implications for Pathogenesis and Therapy in Parkinson’s Disease. Frontiers in Pharmacology 10 (2019).

71. H. Lee, J. Scott, H. Griffiths, J. E. Self, A. Lotery, Oral levodopa rescues retinal morphology and visual function in a murine model of human albinism. Pigment Cell Melanoma Res 32, 657–671 (2019).

72. M. H. Brilliant, K. Vaziri, T. B. Connor, S. G. Schwartz, J. J. Carroll, C. A. McCarty, S. J. Schrodi, S. J. Hebbring, K. S. Kishor, H. W. Flynn, A. A. Moshfeghi, D. M. Moshfeghi, M.E. Fini, B. S. McKay, Mining Retrospective Data for Virtual Prospective Drug Repurposing: L-DOPA and Age-related Macular Degeneration. The American Journal of Medicine 129, 292–298 (2016).

73. T. Mathis, F. Baudin, A.-S. Mariet, S. Augustin, M. Bricout, L. Przegralek, C. Roubeix, É. Benzenine, G. Blot, C. Nous, L. Kodjikian, M. Mauget-Faÿsse, J.-A. Sahel, R. Plevin, C. Zeitz, C. Delarasse, X. Guillonneau, C. Creuzot-Garcher, C. Quantin, S. Hunot, F. Sennlaub, DRD2 activation inhibits choroidal neovascularization in patients with Parkinson’s disease and age-related macular degeneration. J Clin Invest 134, e174199.

74. M. Contin, G. Lopane, L. M. B. Belotti, M. Galletti, P. Cortelli, G. Calandra-Buonaura, Sex Is the Main Determinant of Levodopa Clinical Pharmacokinetics: Evidence from a Large Series of Levodopa Therapeutic Monitoring. Journal of Parkinson’s Disease 12, 2519–2530 (2022).

75. K. C. Chesler, C. T. Motz, K. L. Bales, R. A. Allen, H. K. Vo, M. T. Pardue, Voluntary oral dosing for precise experimental compound delivery in adult rats. Lab Anim 56, 147–156 (2022).

76. D. Deleu, S. Sarre, G. Ebinger, Y. Michotte, The effect of carbidopa on the pharmacokinetics and metabolism of intravenously administered levodopa in blood plasma and skeletal muscle. Naunyn Schmiedebergs Arch Pharmacol 348, 576–581 (1993).

77. A. J. Feola, R. S. Allen, K. C. Chesler, M. T. Pardue, Development of an Automated Electroretinography Analysis Approach. Transl Vis Sci Technol 12, 14 (2023).

78. G. Martin, K. L. Wefelmeyer, F. Bucher, G. Schlunck, H. T. Agostini, Arteriovenous crossing in retinal vessels of mice, rats, and pigs. Mol Vis 26, 731–741 (2020).

79. F. Alonso, L. Li, I. Fremaux, D. P. Reinhardt, E. Génot, Fibrillin-1 Regulates Arteriole Integrity in the Retina. Biomolecules 12 (2022).

80. J. Li, J. Choi, X. Cheng, J. Ma, S. Pema, J. R. Sanes, G. Mardon, B. J. Frankfort, N. M. Tran, Y. Li, R. Chen, Comprehensive single-cell atlas of the mouse retina. iScience 27 (2024).

