## Supplemental Figure 1-3 for "L-DOPA treatment promotes sustained neurovascular and synaptic homeostasis in the diabetic retina"

Eli Chlan, *et al.*

**This PDF file includes:**

Figs. S1 to S3

**Other Supplementary Materials for this manuscript include the following:**

Data S1 to S2

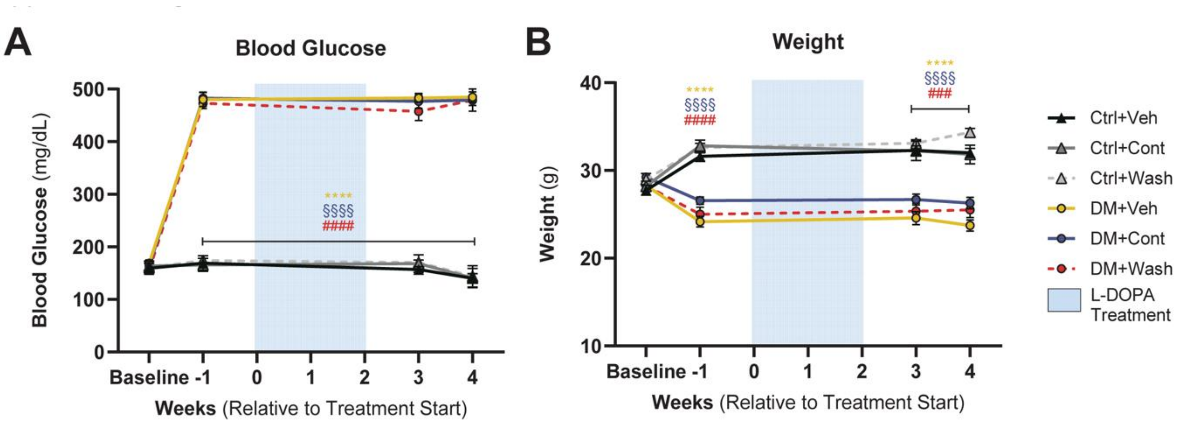


**Fig. S1. Diabetic mice showed consistent blood glucose elevation and lack of weight gain.** (**A**) Blood glucose (mg/dL) was significantly elevated in diabetic mice prior to L-DOPA treatment (-1 wk treatment), as well as after treatment (3-4 week treatment). Control mice did not show BG elevation with vehicle or treatment. (**B**) All diabetic mice maintained significantly lower body weight than control counterparts, with weight consistent within groups across treatment time period. Data shown as mean ± SEM; (*) DM+Veh vs. Ctrl+Veh, (§) Ctrl+Veh vs. DM+Cont, (#) Ctrl+Veh vs. DM+Wash; ***$p < 0.001$,****$p < 0.0001$.

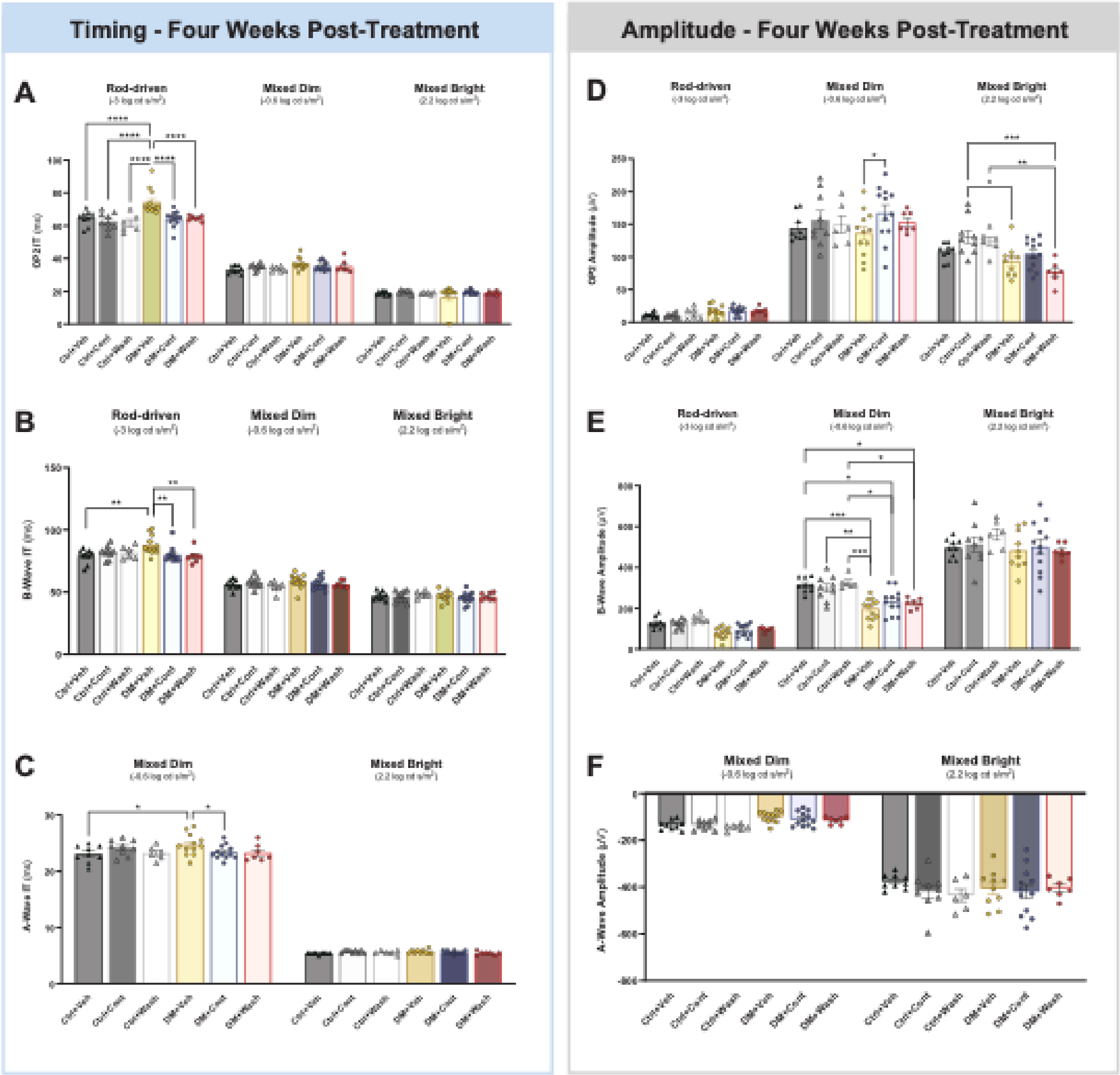


**Fig. S2. Rod-driven oscillatory potential timing was the primary L-DOPA-sensitive ERG component in diabetic mice.** (**A**) Rod-driven oscillatory potential implicit timing was elevated in DM+Veh only. (**B**) Rod-driven b-wave implicit timing showed a similar specificity for delay in DM+Veh mice, but without significant differences compared to Ctrl+Cont and Ctrl+Wash. (**C**) Limited to mixed (rod/cone) dim and mixed bright conditions, mixed dim a-wave timing appeared delayed in DM+Veh mice compared to Ctrl+Veh and DM+Cont mice. (**D**) Oscillatory potential amplitude differences appeared only under mixed dim and mixed bright conditions, without consistent differences between DM+Veh and control mice. (**E**) Mixed dim b-wave amplitude was reduced in DM+Veh mice compared to controls, as well as DM+Cont and DM+Wash mice compared to Ctrl+Veh and Ctrl+Wash. (**F**) No significant differences in A-wave amplitude under mixed dim and bright conditions were seen across diabetic and treatment groups. Data shown as mean ± SEM; *p<0.05, **p<0.01,***$p$ < 0.001,****$p$ < 0.0001.

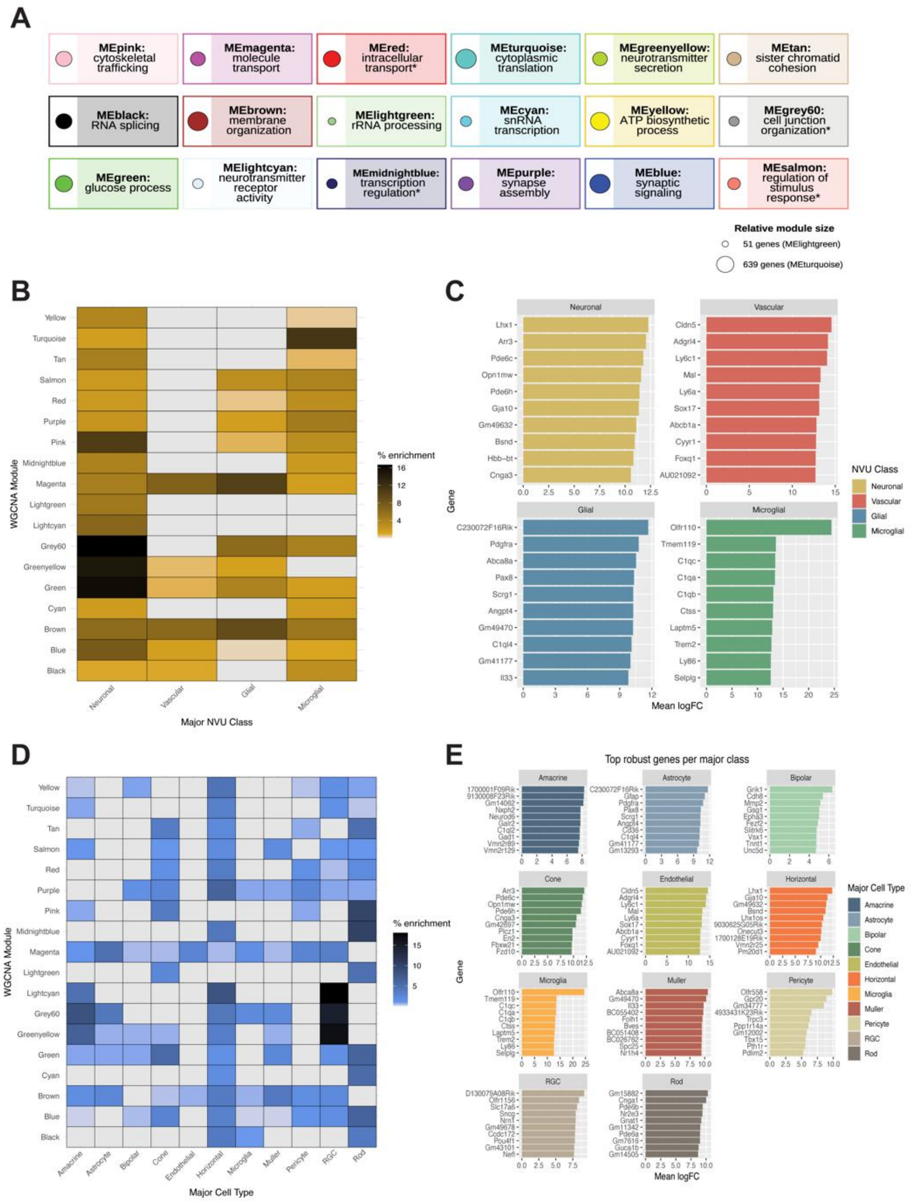


Fig. S3. **Cell-type enrichment and neurovascular unit class enrichment was limited across WGCNA modules.** (**A**) WGCNA modules with summary identity (overarching term from GO:BP) and size (relative circle size). Asterisk indicates modules without significant GO:BP terms. (**B**) Heatmap of major NVU class percent enrichment across WGCNA modules, with a neuronal enrichment across all modules alongside heterogenous spread of vascular, glial, and microglial enrichment. (**C**) Top 10 genes per NVU class, filtered from top 2000 genes unique to each NVU class (logFC > 0.25, padj < 0.05). (**D**) Heatmap of major retinal cell type percent enrichment within WGCNA modules. D) Top 10 genes per major cell class, filtered from top 1000 genes unique to each major retinal cell class.

**Data S1. (separate file) Retinal gene list for identification of major cell class enrichment.**

**Data S2. (separate file) Retinal gene list for identification of neurovascular unit enrichment.**
